# Benzo(a)pyrene (B[a]P) and its adverse effects on male testis

**DOI:** 10.1101/357038

**Authors:** Wei Liu, Aihua Gu

## Abstract

It has been proved that Benzo(a)pyrene (B[a]P) is mutagenic in somatic cells, whereas the adverse effect of BaP on male reproduction remains unclear. To investigate whether it can pass through the blood-testis barrier (BTB) and its potential reproductive toxicology and molecular mechanisms, mice were exposed to B[a]P (there are two doses, that is 13mg/kg body weight and 26 mg/kg body weight; three times per week) during 6 weeks and sacrificed 6 weeks after the final exposure to obtain B[a]P-exposed testis, blood and others. Electron microscopy analysis was performed to confirm whether the integrity of BTB and the ultra-structure changes in testes of B[a]P treated mice, which showed that the integrity of the BTB was disrupted, accompanied with the structure of sertoli cells seriously damaged, including the integrity of the nuclear membrane of the sertoli cells impaired and the basement membrane of the seminiferous tubules disrupted. X-ray imaging in vitro told us that BaP can overgo the BTB and gathered in the testis of mice. We found the significantly decreased expression of ZO-1, occludin, N-cadherin, vimentin and claudin-1 in the testes of B[a]P treated group by immunofluorescence detection. B[a]P induced BTB component protein decreased were also found in TM4 cells exposed to 5μmol/L B[a]P for 24h. We found a significantly decrease of testosterone level and a significantly increase of estrogen level in the serum of treated groups comparing with the control one by radioimmunoassay. TM4 cells, MLTC-1 cells and GC-2 cells was cultured with medium contains B[a]P. MTT Cell Proliferation and Cytotoxicity Assay, cell apoptosis analysis, FACScan analyzer, We observed apparent increase of TM4 and GC-2 cells apoptosis after expose to B[a]P for 24h. B[a]P induced TM4 cell, GC-2 cell and MLTC-1 cell G2/M phase cell arrest. In conclusion, these results suggested that BaP has an adverse impact on male reproduction, it can cross the blood-testis barrier and damage it, the component proteins of the BTB significantly decreased, it can also produce adverse impact on male germ cells.

## 1. Introduction

Benzo(a)pyrene (B[a]P), is a side-product of incomplete combustion or burning of organics (carbon-containing) such as gasoline, cigarettes, and wood [1]. It is generally found in cigarette smoke, in broiled and grilled foods with other PAHs, and as a minor-product of a lot of industrial production processes[1]. BaP is also detected in some water sources, outdoor air, and indoor air[1]. A study of the concentrations of BaP in indoor dustfall, corporal air, ambient air, and food samples was carried out in a study of children’s exposure to PAHs in Minnesota, and found BaP in 43-58% of various kinds of air specimens, 19% of household dust specimens, and 22% of food specimens [2]. As a typical kind of polycyclic aromatic hydrocarbons, BaP is a ubiquitous compound in our everyday life and can jeopardize our healthy. It has been proved to be associated with reduced sperm numbers and abnormal sperm morphology [3, 4]. Both experimental and epidemiologic studies have reported the adverse effects of PAHs and dioxins on male reproductive function. Whereas it is stilled unclear how BaP affects male reproduction, the adverse impact of BaP on male germ cells and whether BaP can overgo the blood-testis barrier.

During spermatogenesis, Sertoli cells supply structural and nutritional support for germ cells, induce phagocytosis of residual bodies, release spermatids, and maintain the spermato-genous microenvironment [5]. What’s more, an important and unique physiological function of Sertoli cells is that they are the main components of the blood-testis barrier (BTB), which protects germ cells from internal antigens and restricts the para-cellular diffusion of various toxic chemicals [5]. Dysfunction of the BTB between Sertoli cells has been considered an important mechanism involved in xenobiotic-induced reproductive toxicity [6]. The BTB is formed by contiguous Sertoli cells near the basal compartment of the seminiferous and constituted of coexisting tight junction (TJ), basal ectoplasmic specialization (ES), and desmosome-like junction[7],it divides the epithelium into basal and adluminal compartments, this division provides a specific microenvironments for spermatogonia and early spermatocytes underneath the BTB[[8, 9].At stage VIII of the seminiferous epithelial cycle, the BTB restructures to allow the pre-leptotene spermatocytes transport from the basal to the apical compartment. This process is regulated by testosterone at the hormone-sensitive stage of the seminiferous epithelium cycle [10, 11]. Therefore, BTB plays an very important role in male reproduction and is associated with male reproductive function. However, no sufficient data have been generated to address the B[a]P-induced disruption of BTB and their role in reproductive toxicity.

In this study, we mainly focused on the male reproductive toxicity of B[a]P, especially it’s adverse effect on the mice BTB and on cells in mice testis. The present results will help reveal the toxicological and physiological importance of B[a]P-induced male reproductive disruption.

## 2. Materials and methods

### 2.1. Animals

All the mice (C57) used in these experiments were bred at the Animal Care Facility of the Nanjing Medical University on a 12L: 12D cycle. Animals had free access to water and a soy-free diet (SDS, Dundee, UK). All experiments involving animals were approved by the Institutional Animal Care and Use Committee of the University of Illinois at Urbana-Champaign and were conducted in accord with the National Institutes of Health Guide for the Care and Use of Laboratory Animal.

### 2.2. Reagents

Benzo(a)pyrene was purchased from J&K. Anti-occludin antibody,anti-claudin-1 antibody, anti-ZO-1 antibody,anti-N-Cadherin antibody, antianti-vimentin antibody anti-ARNT+ARNT2 antibody and anti-AHR antibody were purchased from abcam.The secondary antibody Anti-Rabbit IgG (H+L), F(ab)2 Fragment (Alexa Fluor^®^555 Conjugate) was purchased from Cell Signaling Technology Inc. (Danvers, MA, USA). The HRP conjugated secondary antibody was purchased from Beyotime and the radioimmunoassay kit was purchased from Beijing North Institute Of Biological Technology. Dimethylsulfoxide (DMSO), penicillin-streptomycin, and trypsin were purchased from Sigma-Aldrich (St. Louis, MO, USA). High-glucose Dulbecco’s-modified Eagle’s medium (DMEM),trypsin and fetal bovine serum (FBS) were purchased were purchased from Hyclone(Logan, UT, USA). The MTT used in our viability assay was obtained from Sigma. An annexin V-FITC apoptosis detection kit (BioVision Inc., Mountain View, CA, USA) was used to detect apoptotic activity.

### 2.3. Animal treatments

Male mice, between 8 to 12 weeks old were sub-chronically exposed to B[a]P (there are two doses, 13mg/kg body weight and 26 mg/kg body weight dissolved in 100 μL coin oil; three times per week) by oral gavage during 6 weeks. Unexposed control mice was treated with100 μL coin oil three times per week during 6 weeks. There are eight mice in each group.

### 2.4. Sample collection

Mice were sacrificed 6 weeks after the final exposure by cervical dislocation. One testis was used for electron microscopy while the contra-lateral testis was used for paraffin sections according to methods described elsewhere[12]. Five micrometer tissue sections were then cut and mounted onto glass slides. One epididymis was used for sperm counting whereas the other was used for paraffin sections. Blood samples were collected via sinus orbital puncture using heparinized pulled Pasteur pipettes. Collected blood samples were used to assess the effect of bioa-vailable BaP on systemic concentrations of testosterone and those of estrogen (LH) that regulate testosterone synthesis. Plasma was harvested from each blood sample by centrifugation at 2000 × g at 4 °C for 10 min and stored at −20 °C until assayed for the above mentioned hormones by radioimmunoassay (RIA)[13]

### 2.5. Transmission Electron Microscope

Testes from control and BaP-treated mice were fixed in 2.5% glutaradehyde in 0.1 mol/L phosphate buffer, pH 7.2 1% osmium tetroxide in 0.1 mol/L phosphate buffer.(all reagents from Electron Microscopy Science, Hatfield, PA). Then put the testes into a series concentration of ethanol to dehydration, the testes were embedded in epoxy resin. (Electron Microscopy Science). Thin sections (60 nm) were cut using a Leica Ultra-cut UCT microtome and transferred to 200-l m mesh copper grids, followed by double-staining with uranyl acetate and lead citrate an EM stain apparatus (Leica, ViennaAustria) according to the instructions provided by the manufacturer, using a program with 30 9 stain I (containing uranyl acetate) and 1 9300 stain II (containing lead citrate), both at room temperature. The testicular ultra-structure was examined using a Philips JEM 1010 Transmission Electron Microscope (Philips, Eindhoven, The Netherlands)

### 2.6. Immunofluorescence and Immunohistochemical detection of the BTB proteins

Immunohistochemistry used antibodies of the BTB proteins, such as ZO-1, occludin, N-cadherin, vimentin,claudin-1 and anti-KI67 antibodies to detect the integrity of the BTB and used antibodies of injury markers to investigate the testis trauma. Mounted tissue sections were incubated at a temperature of 60 degrees for 20 minutes. Sections were deparaffinized with a graded series of xylene, and rehydrated in a descending graded alcohol series followed by antigen retrieval with boiling 0.01M citrate buffer (pH 6) by pressure-cooking slides at full pressure for 15 min and then cooling at room temperature. Wash the sections with 0.01M Phosphate Buffered Saline(PBS)for 3 times, 5 minutes per time. Incubating slides in 3% hydrogen peroxide for 10 min at room temperature was used to block endogenous peroxidase activity. Wash steps were carried out as described previously. Nonspecific binding was blocked using 5% bovine serum albumin (BSA)diluted in PBS and incubate at room temperature for 40 min.Sections were incubated with rabbit anti-mouse anti-ZO-1, anti-occludin, anti-N-cadherin, anti-vimentin, anti-claundin and anti-KI67 antibodies diluted 1:100,1:100,1:200,1:700, 1:100 individually in PBS in wet box at 4°C overnight. Wash steps were carried out as described previously followed by incubating sections with the horseradish peroxidase-labeled goat anti-rabbit secondary antibody at 37°C for 60 min for N-Cadherin,claudin-1, vimentin and KI67. Wash steps were carried out as described previously. Then sections were visualized with 3,3′-diaminobenzidine (DAB; Dako)and counterstained with Harris’hematoxylin. At the last, sections were dehydrated and mounted with neutral resins. Slides were photographed using a Provis microscope fitted with a Canon DS126131 camera. For ZO-1 and Occludin, incubating sections with the a fluorescent-labeled secondary antibody at 37°C for 40 min. Wash the sections with 0.01M Phosphate Buffered Saline(PBS)for 3 times, 5 minutes per time. Wipe PBS outside of specimens with filter paper before nuclear counterstain with Dapi for 5 minutes. Wash steps was performed as described above. Buffered glycerol mounting following wipe PBS outside of specimens with filter paper. Using a fluorescence microscope to photograph.

### 2.7. Radioimmunoassay

Plasma samples were analyzed for testosterone and estrogen using RIA following the manufacturer’s instructions. The extraction of testosterone from plasma samples before analysis was one of the pivotal steps in this assay. As a consequence, the total testosterone values will be reported as the result [13]. Samples from animals treated with B[a]P were assayed for testosterone as representative samples. The sensitivity of this assay was 2 pg/tube and the intra-and inter-assay coefficients of variation (CVs) were 3.3% and 6.7%, respectively.

Concentrations of estrogen in plasma samples were determined by the use of RIA kit. Samples from animals treated with B[a]P were assayed for estrogen as representative samples as in the case of testosterone. The sensitivity of this assay was 2 pg/tube and the intra-and inter-assay coefficients of variation (CVs) were 3.3% and 6.7%, respectively.

### 2.8. Cell culture

TM4 mouse Sertoli cells were cultured in DMEM supplemented with 10% FBS and antibiotics (10,000 units/ml penicillin and 10,000 μ g/ml streptomycin sulfate) in a humidified atmosphere of 5% CO2 at 37°C.MLTC-1cells were cultured in 1640 medium. Our toxicity studies were conducted using serum-free DMEM or 1640 medium. The BaP solution was dissolved in DMSO at a concentration of 10mg/ml, this solution was freshly diluted to the appropriate concentration (0.3125 - 20 μ mol/L)with culture medium. Culture medium replenished with DMSO served as the control in each experiment.

### 2.9. Cell viability, cell cycle and cell apoptosis analysis

The viability of TM4 cells,GC-2 cells and MLTC-1 cells were assessed using 3-(4,5-Dimethylthiazol-2-yl)-2,5-diphenyltetrazolium bromide(MTT), cell cycle were analyzed following exposure to 1,3-BP using a FACScan analyzer and cell apoptosis analyzed using dual annexin V-FITC and propidium iodide (PI) staining. Cells were processed according to the methods described elsewhere[14].

### 2.10. Immunocytochemical detection of the Cytotoxic effects of BaP on TM4 cells

Placed sterilized 20mm coverslips into a 6-well plate. Seeding cells at a density of 2*10ˆ4/ml medium into the 6-well plate for cells to climb on the coverslips.Cell contamination was carried out five days later. Cells were exposed to 5μmol/L BaP for 10 hours. Wash the coverslips with 0.01M Phosphate Buffered Saline(PBS)for 3 times, 5 minutes per time followed by fixing in 4% paraformaldehyde for 15 minutes. The coverslips was then drying at room temperature for 5 minutes. Wash steps were performed as described above. Incubating the coverslips with 0.5%Triton X-100 diluted in Dulbecco’s Phosphate Buffered Saline(DPBS) for 20 min at room temperature. Endogenous peroxidase activity used 3% hydrogen peroxide incubating coverslips for 10 min at room temperature to block. Nonspecific binding was blocked using 5% bovine serum albumin (BSA)diluted in DPBS and incubate at at room temperature for 20 min. Coverslips were incubated with rabbit anti-mouse anti-claudin-1,anti-ARNT+ARNT2 antibodies all diluted 1:200 in PBS in wet box at 4°Covernight. Incubating with secondary antibody was processed as described previous. Additionally immunocytochemistry / immunofluorescence used DAPI to stain nuclear for 5 min, mounting sections with buffered glycerol and used a fluorescence microscope to photograph.

### 2.11. Western blot analysis of TM4 cell proteins

Cells were lysed in RIPA buffer (150 mM NaCl, 50 mM Tris, pH 7.4, 1 mM EDTA, and 0.1% NP-40) containing protease inhibitors (20 mg/ml leupeptin, 10 mg/ml pepstatin A, 10 mg/ml chymostatin, 2 mg/ml aprotinin, and 1 mM PMSF). The homogenate was clarified by centrifugation at 15,000 g for15 min and the concentration of protein in homogenates was determined by the Bio-Rad protein assay (Bio-Rad Laboratories, Hercules, CA, USA). Electrophoresis was carried out at 100 V until the dye front reached the bottom of the gel. The proteins were transferred from the gel onto a polyvinylidene difluoride membrane using an electroblotting apparatus according to the manufacturer’ s protocols (Bio-Rad Laboratories). The membranes were incubated in 5% defatted milk powder for 4 h at room temperature to block nonspecific binding. The blocked membranes were incubated with gentle agitation overnight at 4 °C - 8°C with the appropriate primary antibodies (1:1000), followed by incubation with peroxidase-conjugated secondary antibodies (1:2000) for 30 - 60 min at room temperature. The reactive bands were visualized using an enhanced chemiluminescence detection system (Amersham-Pharmacia, Piscataway, NJ, USA) according to the manufacturer’s instructions.

## 3. Results

### 3.1 The effect of B[a]P on mice development

Compared with the control group, there was no significant change in the body weight at the doses of 13mg/kg BW, whereas, we observed a significant decrease in body weight at the doses of 26mg/kg BW(all p<0.05),Fig 1A. After exposing to B[a]P at both 13mg/kg BW and 26mg/kg BW, we found no significant changes of mice heart weight, liver weight, testis weight, in contrast to the control group ( P> 0.05), Fig 1B-D, suggesting that the mice development was not significantly effected by B[a]P.

**Fig.1.**
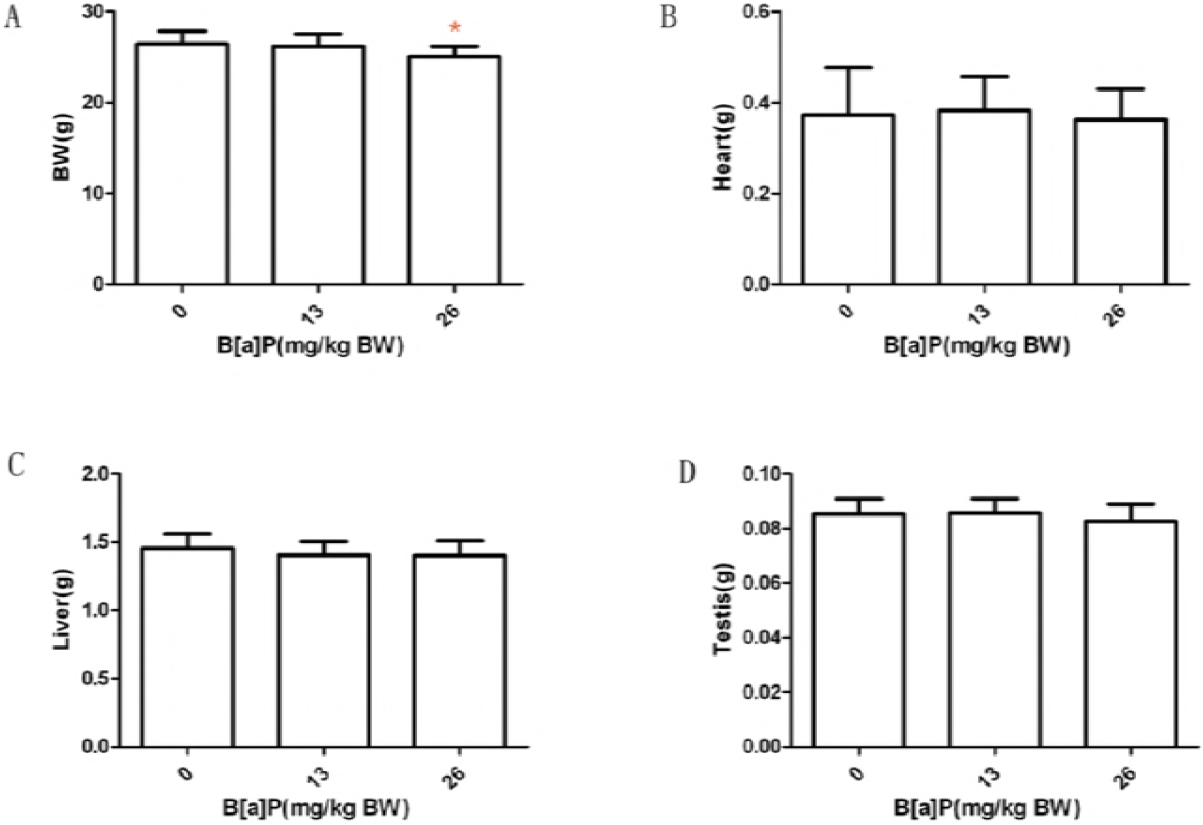
Effects of B[a]P on the mice development. Mice were dosed with either 13mg/kg or 26mg/kg of B[a]P for 6 weeks via gavage and organs such as testis, heart, liver were collected. Mice body weight, testis weight, liver weight and heart weight were evaluated. Mice exposed to 26mg/kg B[a]P showed a significant decrease in body weight (A). Mice heart weight (B), liver weight (C) and testis weight (D) showed no significant change after exposing to B[a]P. Data are mean ± SD (n = 8)

### 3.2 B[a]P-Induced Male Reproductive Damage

B[a]P-Induced Male Reproductive Damage After exposing to B[a]P at doses of 13mg/kg BW and 26mg/kg BW, to confirm this, the testicular morphology was also evaluated. In the control group, the structure and morphology of seminiferous tubules, Sertoli cells, various stages of germ cells, and spermatozoa in the seminiferous epithelium were normal, Fig 2A. However, in the groups treated with B[a]P at doses of 13mg/kg BW and higher, significant increase in Sertoli cell vacuolization and derangement of the cell layers were observed(Figs. 2B). Dislocated immature germ cells were found in the lumens of seminiferous tubules in both group (Fig. 2C).

**Fig.2.**
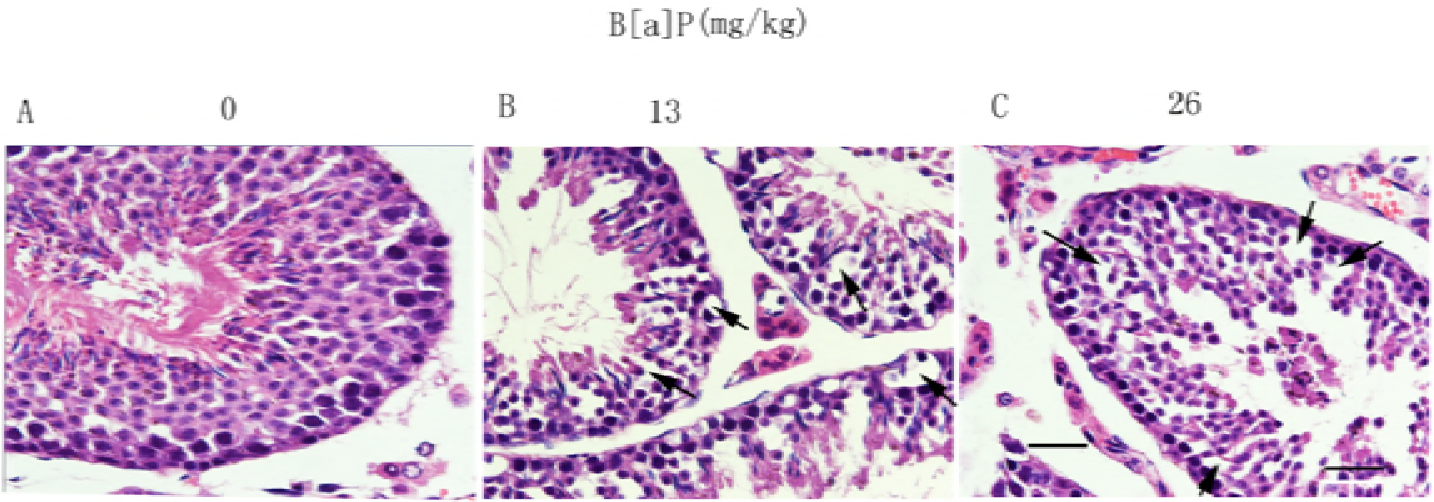
Effects of B[a]P on seminiferous tubules under light microscopy. (A–C) Light microscopy (scale bar = 50 μm). The black arrow indicates Sertoli cell vacuolization.

To confirm these results, the ultra-structure of seminiferous epithelia was evaluated by electron microscopy. The control group showed a normal ultra-structure, with a Sertoli cell nucleus and two germ cell nuclei adjacent to the basement membrane; the cellular layer was clear, and adjacent cells were compact and intact (Figs.3A,a). However, in the groups treated with B[a]P at doses of 13 mg/kg and higher, a significant increase in the vacuolization of Sertoli cells was found (Figs. 3B,b, black asterisk), the basement membrane and nuclear membrane of the Sertoli cells were disrupted(Fig.3C,c).These results indicate that B[a]P damaged seminiferous tubules and specifically disrupted Sertoli cells.

**Fig.3.**
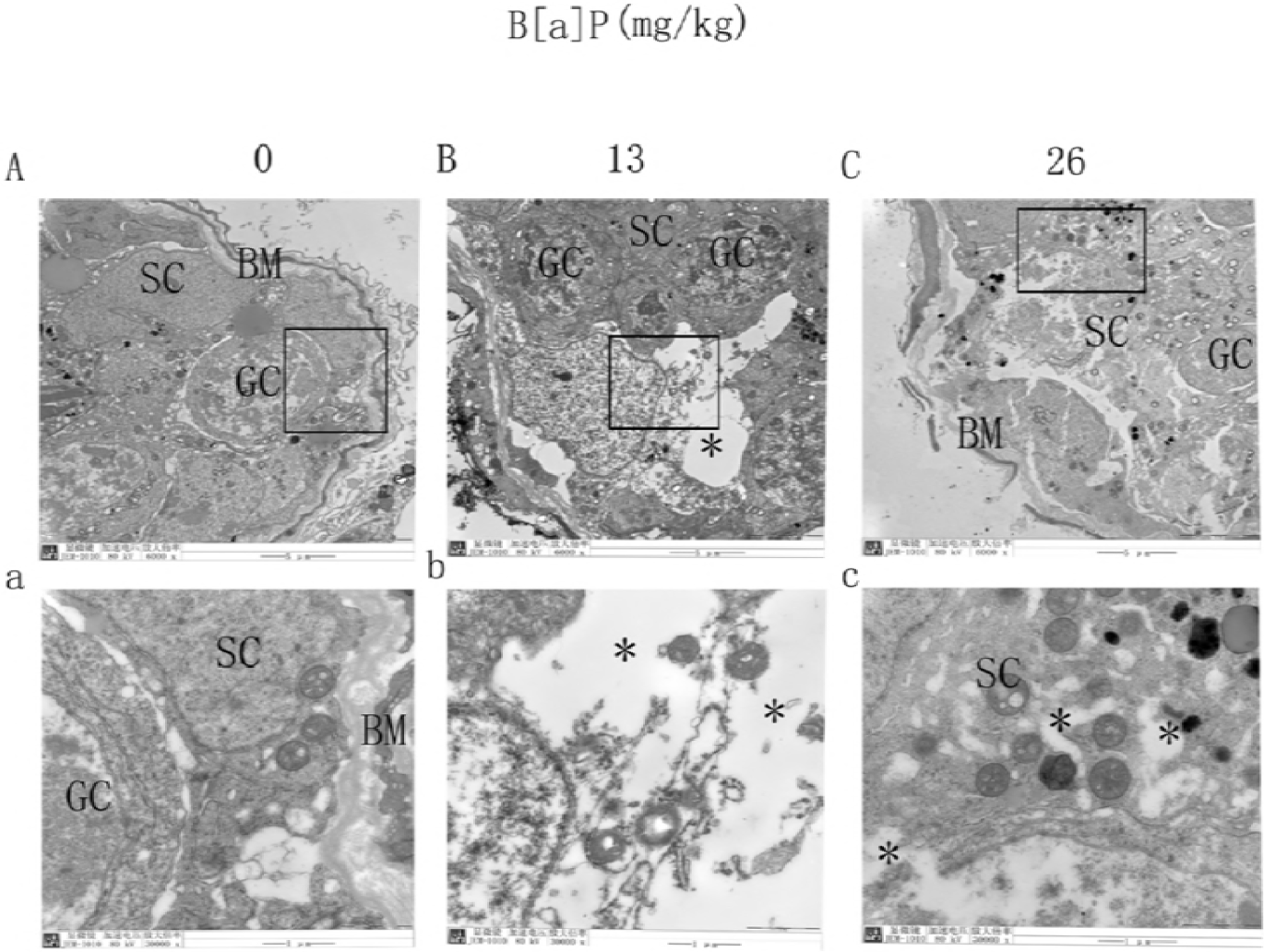
Effects of B[a]P on seminiferous tubules under electron microscopy. (A-C) Electron microscopy (scale bar = 5 μm). (a–c) Magnification of boxed areas from A–C (scale bar = 1 μm). SC, Sertoli cell; GC, germ cell; BM, basement membrane.The black asterisk indicates Sertoli cell vacuolization and vacuoles. *p < 0.05 compared with control.

### 3.3 B[a]P had a significant impact in mice endocrine function

Expose to B[a]P (13mg/kg and 26mg/kg body weight) depressed total plasma testosterone (P< 0.05), Fig 4A. Expose to B[a]P at 13mg/kg BW had no significant influence on total plasma estrogen level ( P>0.05); Expose to B[a]P at 26mg/kg BW significantly increased total plasma estrogen concentrations compared with controls ( P< 0.05),Fig. 4B. These results told us that B[a]P had a significant impact in mice endocrine function.

**Fig.4.**
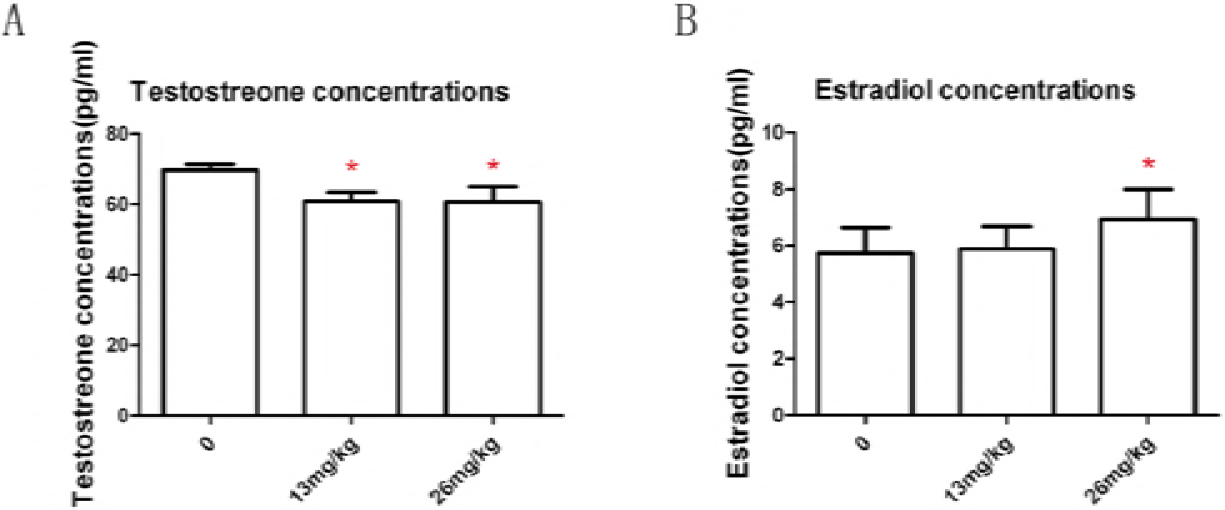
Effect of B[a]P treatment on testosterone concentrations in serum (A) and estradiol concentrations in serum (B). Testosterone and estradiol levels were measured by RIA. Data are mean ± SD (n = 8). *p < 0.05 compared with control.

### 3.4. Effects of BaP on cell viability

To investigate whether or not BaP effect cell viability, TM4 cells, GC-2 cells and MLTC-1 cells were exposed to different concentrations of BaP (0.3125-20 μ mol/L) for 24 h and cell viability was assessed by MTT assay. As described in Fig. 5A,TM4 cells viability were not significantly altered after concentrations of BaP ranging between 0.3125 μ mol/L and 20 μ mol/L exposure for 24h. GC-2 cells viability were significantly altered after concentrations of BaP at 10 μ mol/L exposure for 24h., Fig. 5B. MLTC-1 cells viability were significantly altered after concentrations of BaP ranging between 2.5 μ mol/L and 20 μ mol/L exposure for 24h, Fig. 5C.

**Fig.5.**
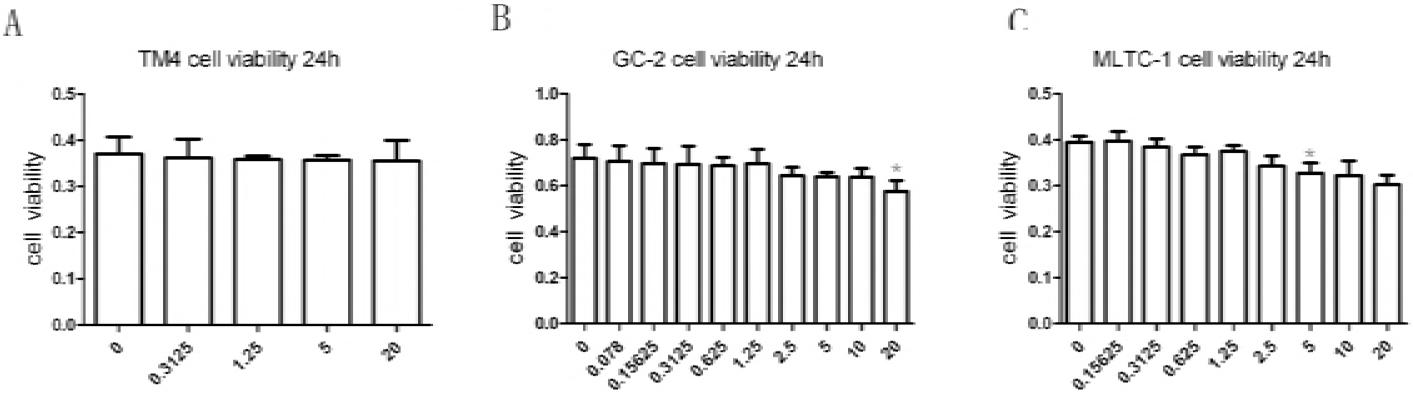
Effects of B[a]P on TM4 cell viability (A), on GC-2 cell viability (B), on MLTC-1 cell viability (C). TM4 cells were treated with 0, 0.3125, 1.25,5, or 20μmol/L B[a]P for 24h. GC-2 cells were treated with 0, 0.078,0.15625,0.3125,0.625,1.25,2.5,5,5,10 or 20μmol/L B[a]P for 24h. MLTC-1 cells were treated with 0, 0.15625,0.3125,0.625,1.25,2.5,5,5,10 or 20μmol/L B[a]P for 24h. At the end of the incubation period,cell viability was estimated by a formazan assay using MTT reagents. The results are from three experiments. Data are mean ± SD (n = 8). *p < 0.05 compared with control.

### 3.5 BaP-induced apoptotic cell death

To determine whether BaP exposure induced apoptotic cell death, fluorescence analysis was then carried out using annexin V-FITC/PI after treating cells with BaP at doses of 0, o.3125 and 5 μ mol/L for 24 h. Apoptosis was observed by fluorescence microscopy and flow cytometry dot plot analysis. TM4 cells exposed to 0.625, 2.5 and 10 μ mol/L BaP for 24 h, comparing to control cells treated with DMSO, Fig. 6A. GC-2 cells exposed to 0.625, 2.5 and 10 μ mol/L BaP for 24 h, comparing to control cells treated with DMSO was shown in Fig. 6B. MLTC-1 cells exposed to 0. 3125 and 1.25 μ mol/L BaP for 24 h, comparing to control cells treated with DMSO was shown in Fig. 6C. B[a]P induced TM4 cells, GC-2 cells apoptosis and it showed no significant influence on MLTC-1 cells.

**Fig.6.**
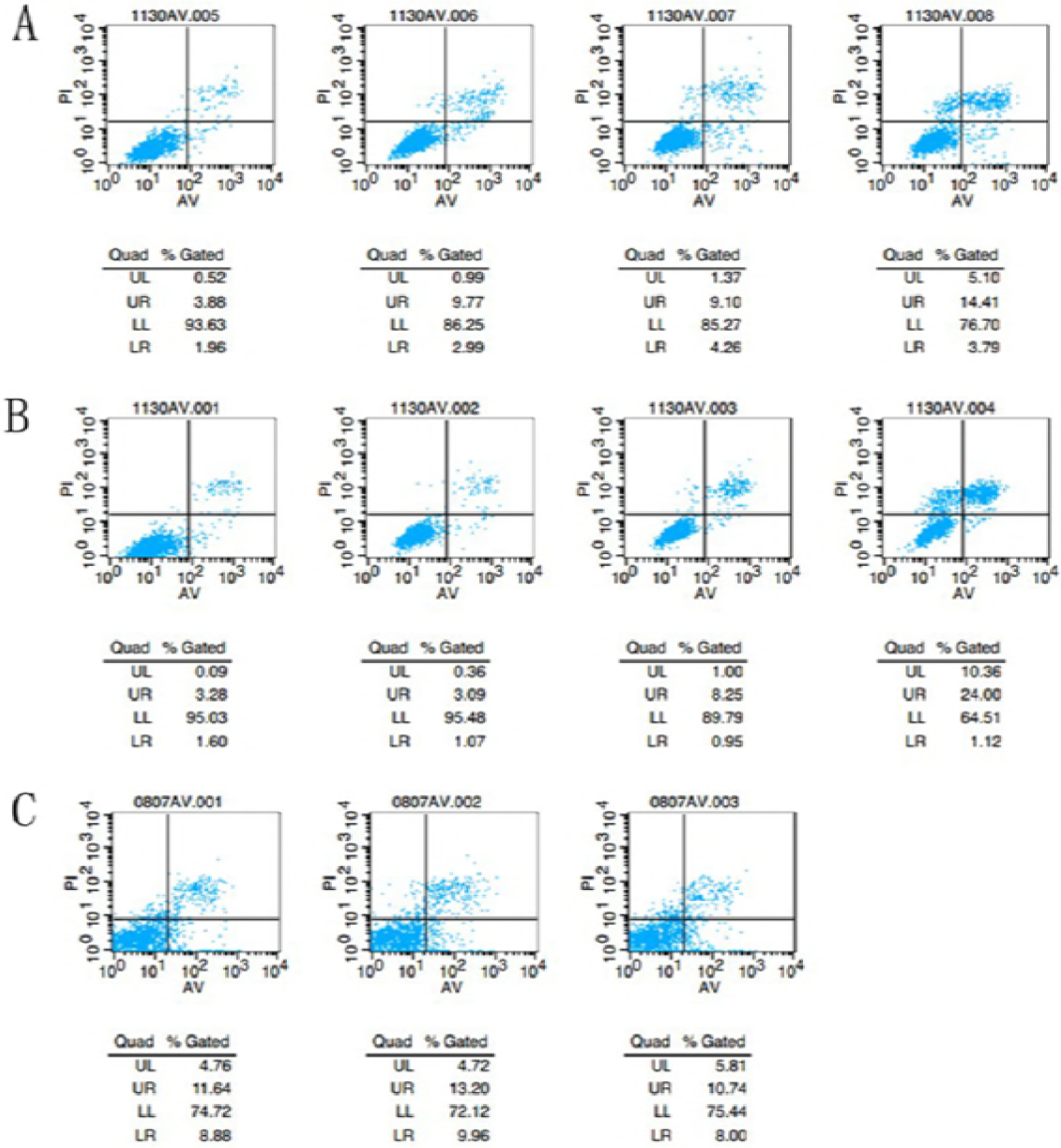
Effect of B[a]P on apoptotsis. Flow cytometry dot plots analysis of annexin-V/PI-labeled TM4 cells (A), GC-2 cells (B), MLTC-1 cells (C). TM4 cells and GC-2 cells were treated with 0.1% DMSO (control) or with 0.625, 2.5, or 10μmol/L B[a]P for 24h, MLTC-1 cells were treated with 0.1% DMSO (control) or with 0.3125 or 1.25μ mol/L B[a]P for 24h. The number in the corner of each quadrant is the percentage of total cells that are detected in that quadrant. Lower left quadrant: viable annexin-V-negative cells; lower right quadrant: annexin-V-positive cells; upper left quadrant: necrotic PI-positive cells; upper right quadrant: late apoptotic or necrotic cells.

### 3.6. BaP induced abnormal cell cycle

To detect wether BaP exposure can change the cell cycle, we used flow cytometry to examine the cell cycle phase distribution. 5 μ mol/L BaP induced TM4 cell G2/M phase cell arrest, Fig. 7A; 5 μ mol/L BaP induced GC-2 cell G2/M phase cell arrest, Fig. 7B; 1.25 μ mol/L BaP induced MLTC-1 cell G2/M phase cell arrest, Fig. 7C.

**Fig.7.**
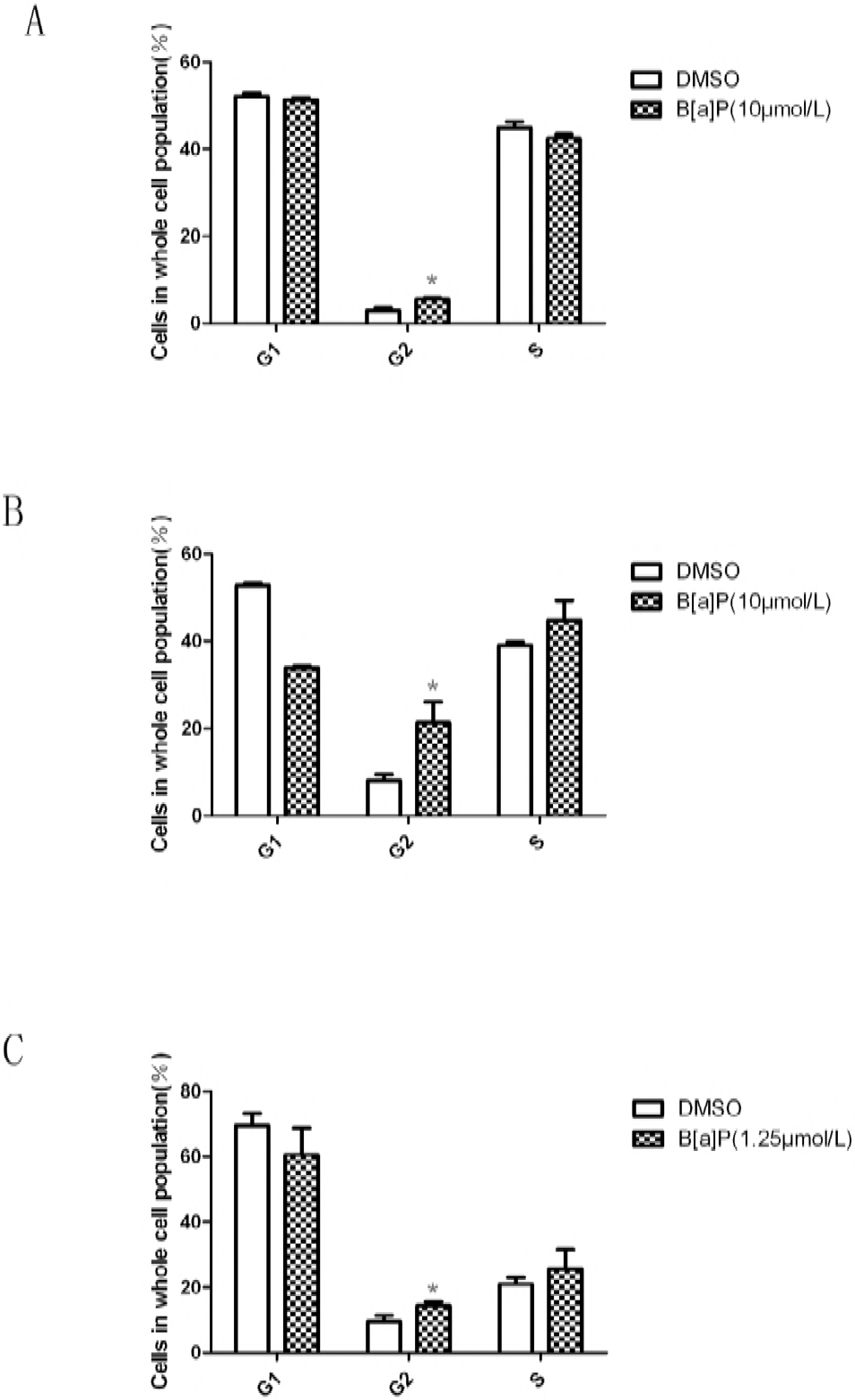
Effect of B[a]P on the cell cycle distribution in mouse TM4 Sertoli cells, GC-2 cells and MLTC-1 cells. Cells were plated in 60-mm dishes at 1.8×10^5^/dish and incubated overnight in an incubator. On the following day, the TM4 cells and GC-2 cells were treated with 10μmol/L B[a]P and incubated for 24h, the MLTC-1 cells were treated with 1.25μmol/L B[a]P and incubated for 24h.The cell cycle distribution was measured by FACScan flow cytometry as described in Section2. The data are expressed as the mean±SD. (*) indicates P<0.05 compared to the untreated controls.

### 3.7. BaP-Induced Structural Changes in the BTB Between Sertoli Cells

Transmission electron microscopy was carried out to detect the adverse effect of BaP on the ultra-structure of BTB. The normal and clear ultrastructure of the BTB between two adjacent Sertoli cells (Fig. 8A) showed a classic basal ectoplasmic specialization structure, actin filament bundles sandwiched between the endoplasmic reticulum membrane and opposing plasma membranes of two adjacent Sertoli cells, and a TJ (white arrow) in the BTB (Fig. 8a). However, significant changes, such as disassembly of the TJ and vacuoles (black asterisk), were observed in the groups treated with BaP at doses of 13 mg/kg and higher (Figs. 8B-D, b-d). These results confirmed the disruption of BTB integrity and function via damage to the ultra-structure of the BTB between Sertoli cells.

**Fig.8.**
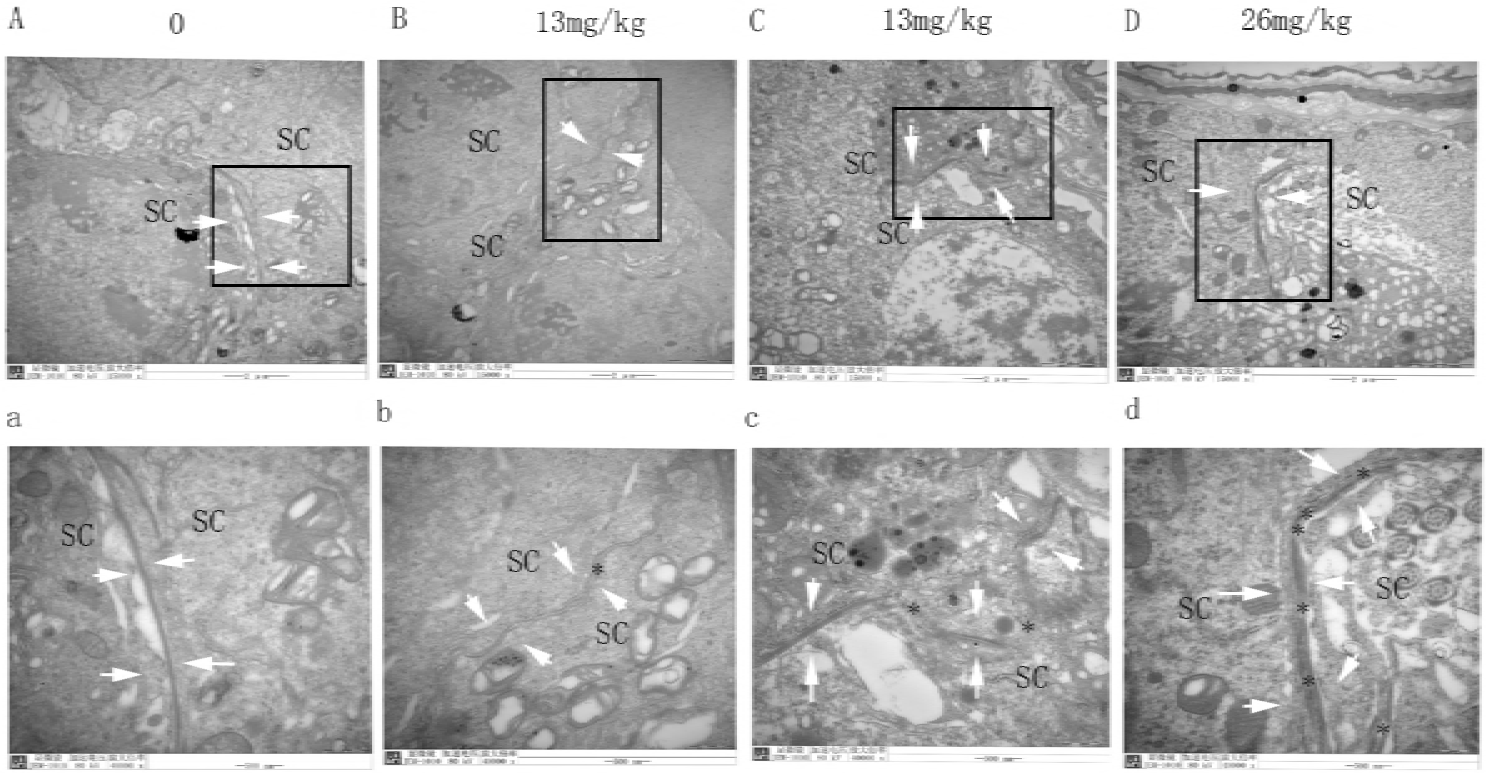
Effects of B[a]P on ultra-structure of the BTB. (A–D)Typified fields (scale bar = 2 μm). The BTB is indicated by opposing white arrow. SC, Sertoli cell. (a–d) Magnification of boxed areas from A–E (scale bar = 500 nm). The black asterisk indicates disassembly of the TJ and vacuoles in a–d.

### 3.8. BaP-Induced Changes in the Expression and Localization of Testicular TJ and GJ Proteins

To investigate the molecular mechanisms of BaP affect the the BTB integrity, gene chips, sections immunohistochemistry was performed to research them.

The change of the BTB functional junction proteins was evaluated in the different treatment groups using immunohistochemistry and immunofluorescence. Occludin, as a tight junction protein, has been detected at the site of the BTB [8]. ZO-1 is a tight junction adaptor protein. Immunofluorescence staining for ZO-1 and occludin was undertaken to assess the state of the tight junctions at the BTB. In control testes, ZO-1 and occludin clearly localized at the site of the BTB, parallel to the basement membrane of seminiferous tubules(Fig.9A,D).After exposed to BaP(13mg/kg), ZO-1 and occludin significantly decreased from the seminiferous tubules(Fig.9B,E)and following high-dose (26mg/kg) BaP treatment, ZO-1 and occludin appeared to be absent from the STs (Fig.9C,F).

**Fig.9.**
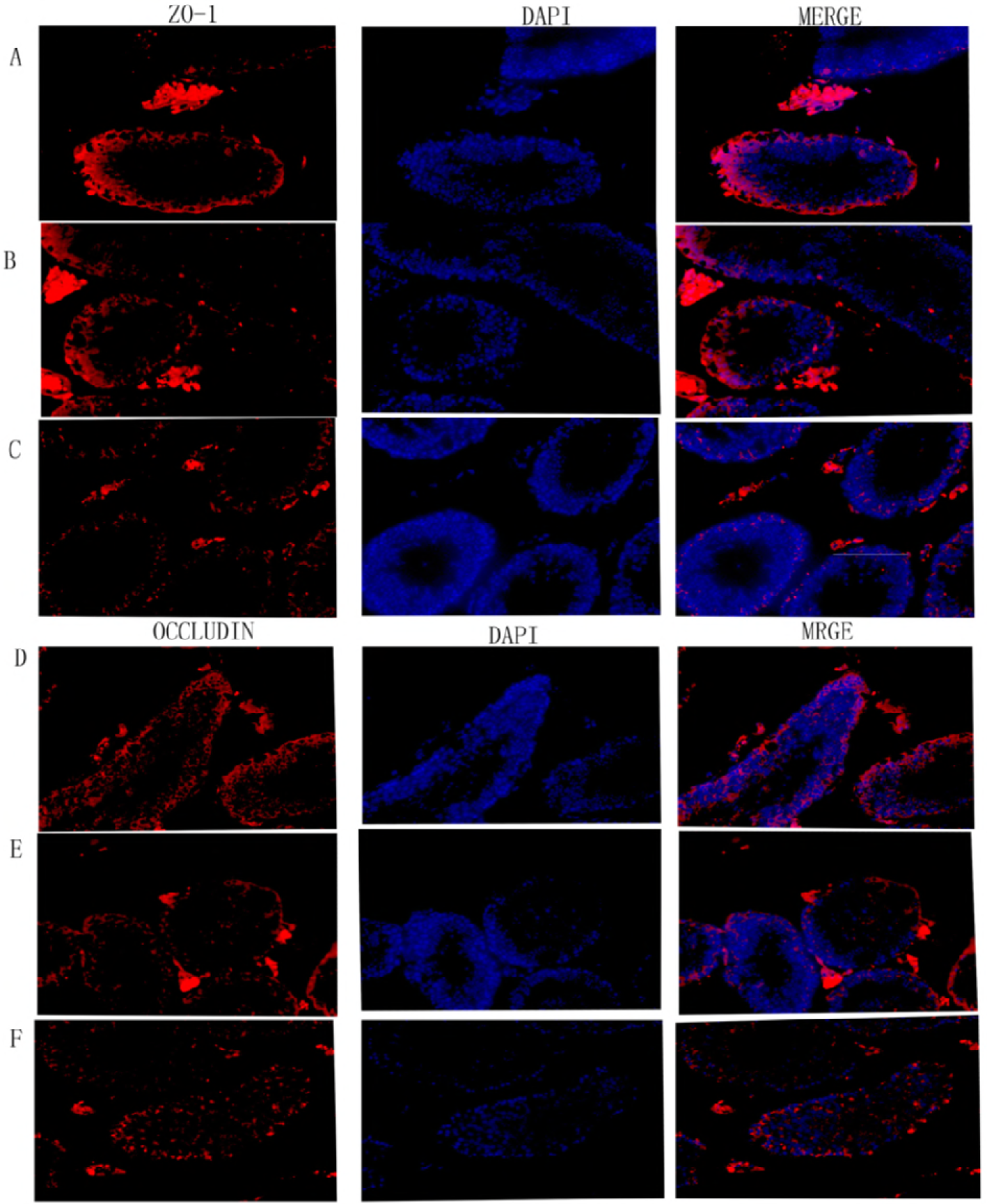
Effects of B[a]P on the expression of proteins related to BTB function. Mice were dosed with either a low or high dose of B[a]P and testes collected 6 weeks later and evaluated for immunoexpression of the BTB function related proteins TJ proteins: ZO-1 (A-C) and occluding (D-F) (red); counterstaining used DAPI (blue). Scale bar represents 100μm. (A,D) Testes from vehicle-treated mice show normal spermatogenesis and the expression of ZO-1 and occludin was normal. (B-C, E-F) Testes from B[a]P treated mice show toxicant-induced damage to spermatogenesis, and the expression of ZO-1 and occludin was decreased.

To assess the condition of adherens junctions in the BTB, we used immunohistochemistry for the vimentin. Vimentin is expressed in the cytoplasm near the abluminal compartment of the sertoli cells (Fig. 10A). After exposed to BaP(13mg/kg), vimentin significantly decreased from the seminiferous tubules(Fig.10B)and following high-dose (26mg/kg) BaP treatment, vimentin appeared to be absent from the STs (Fig.10C). These results expounded that treatment with 13 mg/kg and 26mg/kg BaP has disrupted the tight junctions and adherens junctions at the BTB.

**Fig.10.**
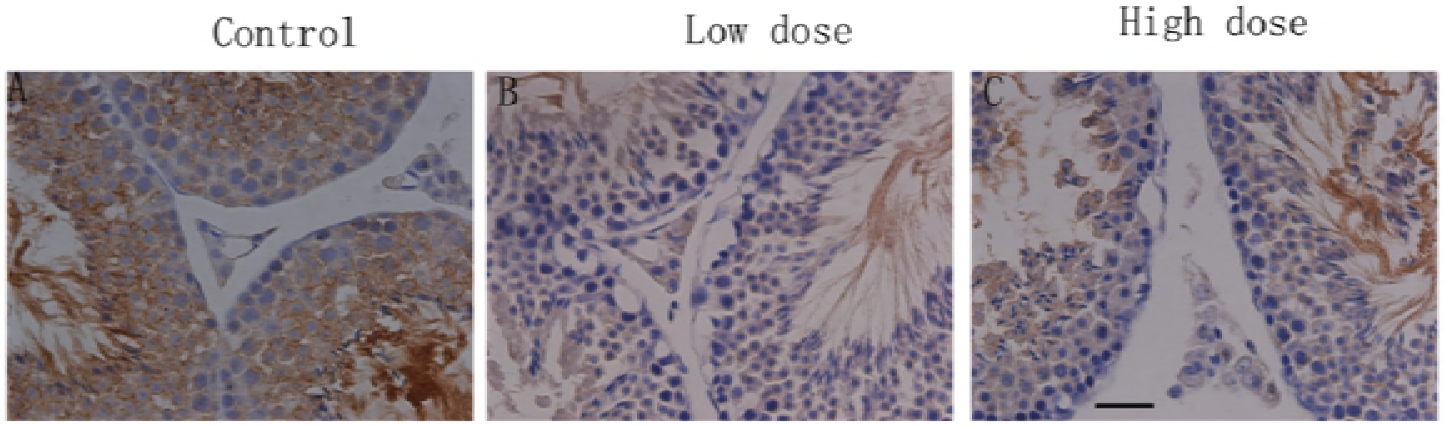
Histological analysis of the effects of treatment with B[a]P on mice testes. Mice were dosed with either a low or high dose of B[a]P and testes collected 6 weeks later and evaluated for immunoexpression of the BTB function related proteins: vimentin (brown); counterstaining used hematoxylin (blue). Scale bar represents 50μm. (A) Testes from vehicle-treated rats show normal spermatogenesis and the presence of vimentin in the cytoplasm near the abluminal compartment of the sertoli cells. (B-C) Testes from 13mg/kg and 26mg/kg B[a]P treated mice show toxicant-induced damage to spermatogenesis, and the expression of vimentin was decreased.

In vitro study, we used western blot and Immunocytochemistry in TM4 cells exposing to BaP to test and verify the result of sections immunohistochemistry and immunofluorescence. In the control slides, claudin-1 was expressed in the cell membrane of the TM4 cells as we can see from Fig.11A. Following cultured with BaP at a dose of 5μmol/L for 24 hours, the expression of claudin-1 was significantly decreased(Fig.11B).The result suggested that the component proteins of BTB were disrupted following BaP exposure.Westernblot told us that the expression of the BTB functional junction proteins in significantly decreased TM4 cells, Fig 12.

**Fig.11.**
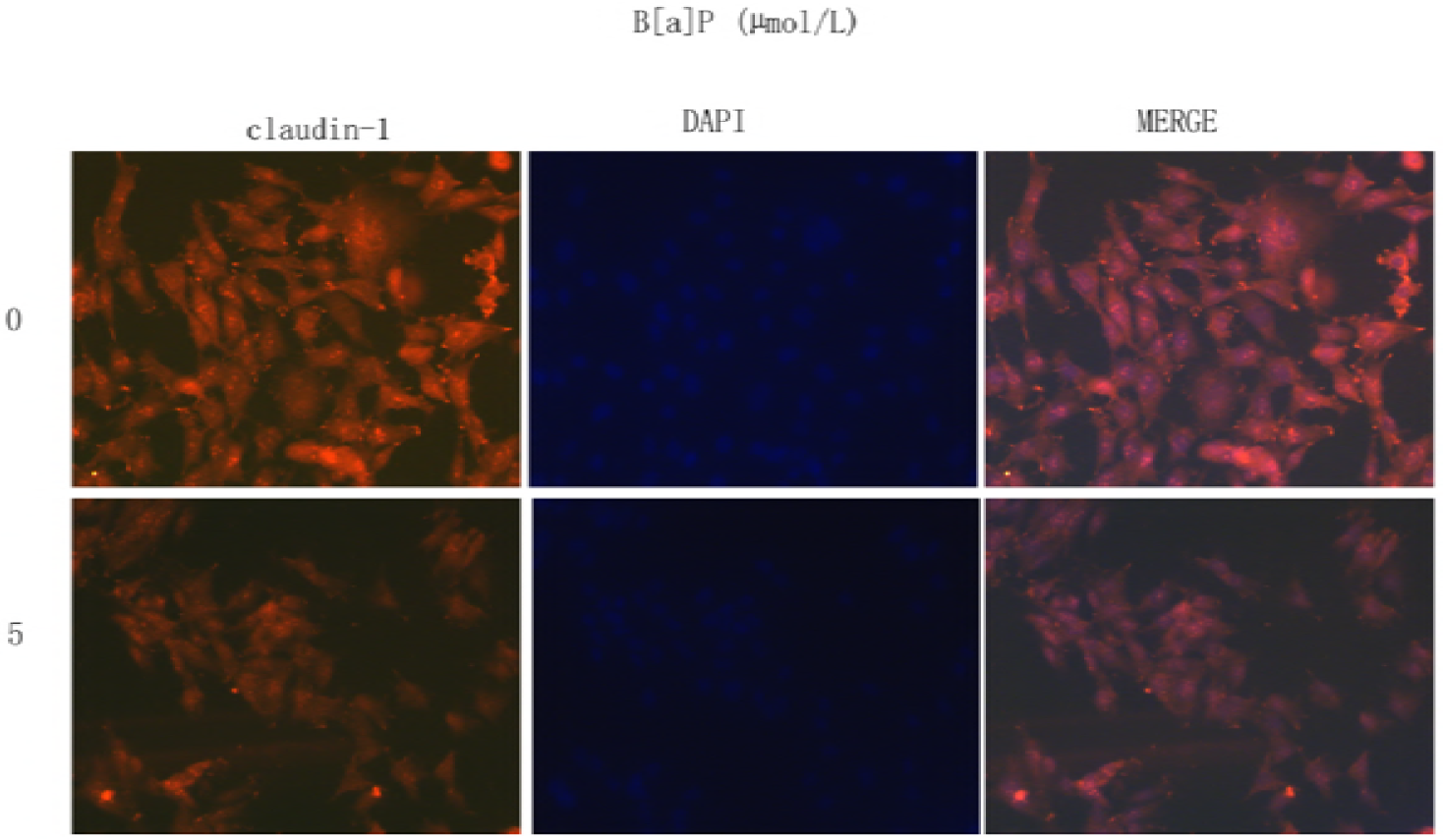
Immunocytochemistry analysis of the effects of treatment with B[a]P on the BTB function related proteins claudin-1 (red) in TM4 cells; counterstaining used DAPI (blue).. TM4 cells were treated with 5μmol/L B[a]P for 24h. (A) In the control slides, claudin-1 was expressed in the cell membrane of the TM4 cells. (B) Following cultured with BaP at a dose of 5μmol/L for 24 hours, the expression of claudin-1 was significantly decreased.

**Fig.12.**
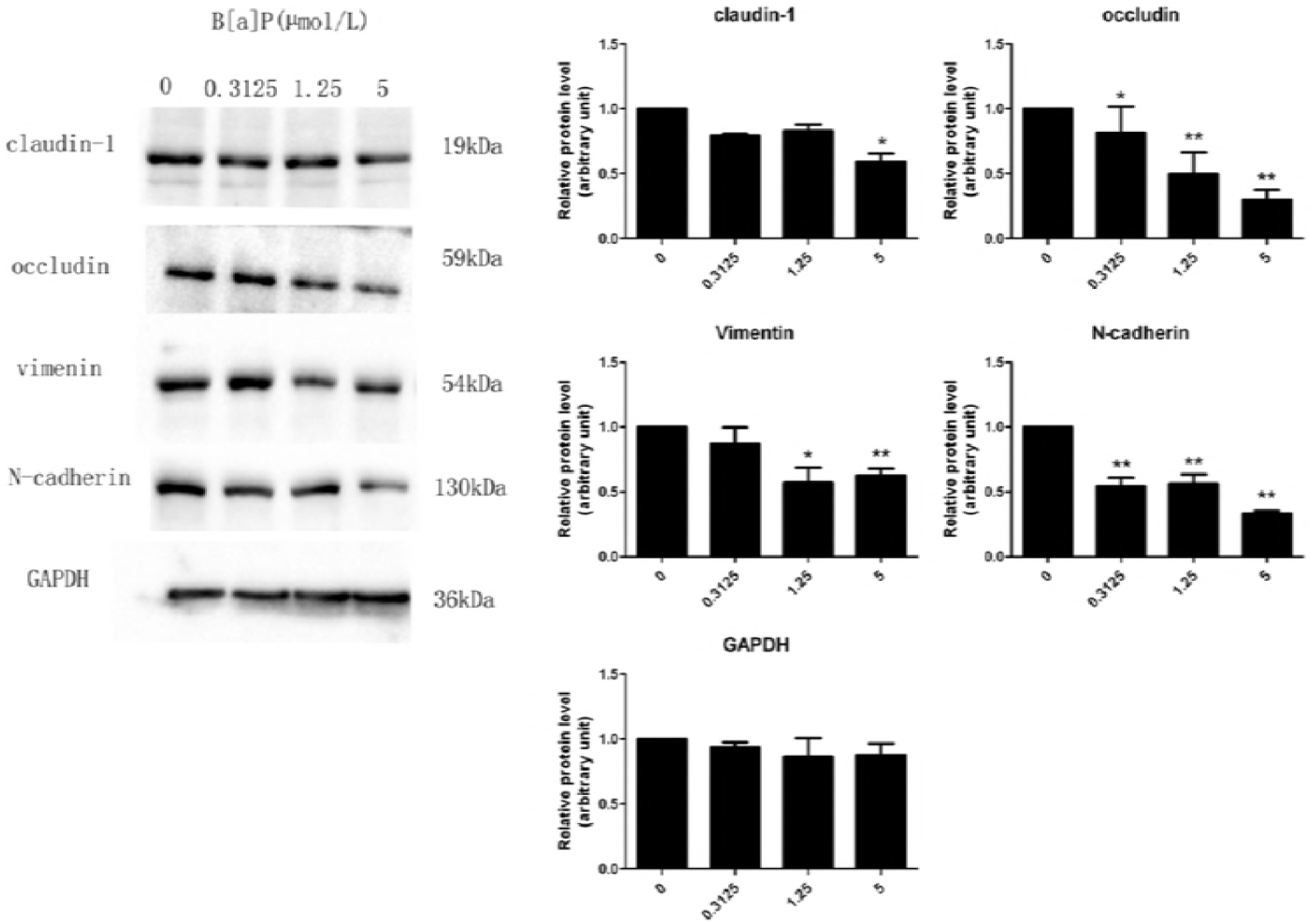
Effects of B[a]P on the expression of proteins related to BTB function in TM4 cells. TJ proteins: ZO-1, claudin-1, and occludin. Basal ectoplasmic specialization proteins: N-cadherin. And vimentin. Each protein (50 μg) was determined by immunoblot using specific antibodies. ImageJ software was used to quantify the integrated band intensity, which was transferred to relative values using corresponding glyceraldehyde-3-phosphate dehydrogenase as an internal reference. The data are expressed as the mean ± SD of three separate experiments from each mouse. *p < 0.05, **p < 0.01, compared with control group.

## 4. Discussion

The BTB is formed by contiguous Sertoli cells near the basal compartment of the seminiferous and constituted of coexisting tight junction (TJ), basal ectoplasmic specialization (ES), and desmosome-like junction[7].The BTB also divides the seminiferous epithelium into the basal and adluminal (apical) compartment so that spermiogenesis can take place in a specialized microenvironment in the apical compartment behind the BTB[15] and it contributes to the immune privilege status of the testis at least in part, so that anti-sperm antibodies are not developed against antigens which express fugaciously during spermatogenesis[15]. In adult mammals, such as rodents and humans, the blood-testis barrier (BTB) is an important and unique ultrastructure in the seminiferous epithelium which maintains spermatogenesis[7].In the rat testis, the BTB awards the ‘railings’ function that supervises the paracellar inflow of water, electrolytes, hormones, and biomolecules between adjacent Sertoli cells and maintains cell polarity[16] and serves as a ‘gatekeeper’ to keep harmful substances from reaching developing germ cells, most notably post-meiotic spermatids[15].Herein, the BTB plays a very important role in male reproduction and is crucial for spermatogenesis and fertility[17].It has been reported that cadmium and bisphenol A exert their adverse effects in the testis by distressing blood-testis barrier function, which successively affects germ cell adhesion in the seminiferous epithelium[17]. Recent studies have also demonstrated that men who exposed to Cd and/or other environmental toxicants have a reduced male fertility, such as reduced sperm count and poor semen quality[18]. Adjudin, a derivative of 1H-indazole-3-carboxylic acid, exerting its effects by disrupting adhesion of germ cells, most notably spermatids to the Sertoli cells locally in the apical compartment of the seminiferous epithelium, behind the blood-testis barrier, thereby inducing release of immature spermatids from the epithelium that leads to infertility[19].In conclusion, xenobiotics can disrupt the BTB and induce male infertility.

Here we studied the influence of BaP on male reproduction, especially the integrity of the BTB, for the first time. Testicular electron microscopy, immunofluorescent evaluation of BTB component proteins (occludin, ZO-1) and immunohistochemistry evaluation of BTB component proteins (vimentin, N-cadherin) were carried out to the present treatment groups to identify whether the integrity of BTB was disrupted following BaP exposure. The present studies show unequivocally that BaP, at doses of 13mg/kg and 26mg/kg, disrupted the BTB as evidenced by testicular electron microscopy, immunofluorescent and immunohistochemistry. The electron microscopy of testis exposed to BaP at both doses showed that the integrity of BTB was disrupted, the ultra-structure of the sertoli cells and spermatogenic cells were damaged seriously, endolysis, abnormality of organelles, even the basement membrane of seminiferous tubule and nuclear membrane of sertoli cells were disrupted compared with the control group. Immunofluorescent and immunohistochemistry found that there was significantly loss of component proteins (occludin, ZO-1, N-cadherin, and vimentin) after exposing to BaP at doses of 13mg/kg and 26mg/kg compared with the control sections. Immunocytochemistry confirmed the results and showed a significantly decrease of the expression of claudin-1 of TM4 cells following exposed to BaP at a dose of 5μmol/L for 24 hours. Above all, the integrity of the BTB of mice was disrupted by BaP exposure. BaP and/or its metabolites disturb the androgen-dependent courses by acting as anti-androgens in target tissues[13].Tight junctions which formes between the Sertoli cells are a major component of the blood-testis barrier and are regulated by testosterone[20]. Testosterone promotes the integrity of the blood-testis barrier in vivo[20, 21]. It has been reported that the tight junctional components and function was disrupted following depletion of testosterone action on Sertoli cells[22].In addition to the androgen, estrogens are also vital for male reproductive function[13]. Herein, we detected the testosterone and estrogen level of mice plasma and found that BaP induced a significantly decrease at doses of 13mg/kg and higher, while the estrogen level significantly increased at dose of 26mg/kg.

The cytotoxicity of BaP on the GC-2 cells which is an immortalized mouse pachytene spermatocyte-derived cell line[23] suggested that BaP exposure may induce genotoxicity in adult men. It has been reported that Leydig cells extensively metabolize BaP[24]. Studies have shown that BaP is extensively metabolized in the male reproductive system[13] The metabolites of exposed BaP may have significantly decreased the population of Leydig cells, thereby, leading to decreased testosterone levels [13]. The change of testosterone and estrogen in testes tumor Leydig cell lines MLTC-1 cells cultured with BaP in our study is consistent with the previously report of subacute exposure to inhaled benzo(a)pyrene (BaP) in Fisher 344 rats.

In order to further investigate the adverse effect of bioavailable BaP on the testis microenvironment of mice where spermatogenesis take place, we focused on the impact of BaP on TM4 cells, MLTC-1 cells and GC-2 cells. The present studies show unequivocally that BaP can induced TM4 cells and GC-2 cells apoptosis and induced TM4 cells, MLTC-1 cells and GC-2 cells G2/M phase cell arrest. BaP also had a adverse effect on the ultra-structure of the male mice testis, a significant increase in the vacuolization of Sertoli cells was found, the basement membrane and nuclear membrane of the Sertoli cells were disrupted.

In summary, the findings of our study strongly indicate for the first time that sub-chronic exposure to bio-available BaP can seriously disrupt the blood-testis barrier in mice with the ultra-structure of the BTB damaged and the BTB component proteins were significantly decreased. In addition, BaP can adversely effect male mice reproduction via inducing suppression of mice plasma testosterone and increased mice plasma estradiol which are involved in BTB regulation, damaging the ultra-structure of sertoli cells in mice testis and changing the normal morphology of the seminiferous epithelium. Furthermore, BaP exerted its adverse effect on the testis microenvironment of mice where spermatogenesis take place and induced abnormal cell cycle and apoptosis of the TM4 cells and GC-2 cells. The long-term effect of BaP treatment on the structural and functional integrity of the blood-testis barrier and male reproduction toxicity outcomes are expected.

